# A non-conserved histidine residue on KRAS drives paralog selectivity of the KRAS G12D inhibitor MRTX1133

**DOI:** 10.1101/2022.12.16.520846

**Authors:** Miles A. Keats, John J. W. Han, Yeon-Hwa Lee, Chih-Shia Lee, Ji Luo

## Abstract

The discovery of small molecule inhibitors against mutant KRAS protein was a recent breakthrough in targeted therapy. Understanding the mechanism of selectivity by KRAS inhibitors and the mechanism of resistance to them can guide the future development of KRAS inhibitors. MRTX1133 was the first non-covalent inhibitor against the KRAS G12D mutant that demonstrated specificity and potency in pre-clinical tumor models. We used isogenic cell lines expressing a single RAS allele to evaluate the selectivity of this compound. We found that in addition to KRAS G12D, MRTX1133 shows significant activity against several other KRAS mutants as well as wildtype KRAS protein. In contrast, MRTX1133 exhibits no activity against both G12D and wildtype forms of HRAS and NRAS proteins. Functional analysis revealed that the selectivity of MRTX1133 towards KRAS is associated with its binding to histidine 95 on KRAS, a residue that is not conserved in HRAS and NRAS. Reciprocal mutation of amino acid 95 among the three RAS paralogs resulted in reciprocal change in their sensitivity towards MRTX1133. Thus, histidine 95 is an essential selectivity handle for MRTX1133 towards KRAS. This knowledge could aid the development of future KRAS-selective inhibitors.

## Main Text

The discovery^1-4^ and clinical development^5,6^ of mutant-selective inhibitors against the small GTPase KRAS has been a recent breakthrough in targeted therapy. All three members of the RAS family of small GTPases, *KRAS, HRAS*, and *NRAS*, are major oncogenes in human cancer. Oncogenic mutation in RAS protein attenuates its GTPase activity and constitutively increases the level of GTP-bound, active RAS in the cell. The MAP kinase (MAPK) pathway, which consists of the RAF/MEK/ERK kinase cascade, is a major downstream effector pathway of RAS that mediates its mitogenic effect^7^. The Glycine 12 to cysteine (G12C) mutant from of KRAS was the first to be successfully targeted by small molecule covalent inhibitors. KRAS G12C inhibitors rely on their ability to cross-link to the mutant C12 residue on KRAS achieve their exquisite selectivity. Two KRAS G12C inhibitors, sotorasib (AMG 510) and adagrasib (MRTX849), were recently approved by the FDA for treating non-small cell lung cancer with KRAS G12C mutation^8^. Recently, MRTX1133 was disclosed as the first non-covalent inhibitor with selectivity for the KRAS glycine 12 to aspartic acid (G12D) mutant. MRTX1133 was evolved from the scaffold of adagrasib through extensive structural optimization to improve its affinity for the KRAS switch II pocket and for selectivity towards the mutant D12 residue on KRAS^9^. The discovery of MRTX1133 demonstrates that KRAS mutant without a cross-linkable side chain can be targeted with conventional, non-covalent inhibitors.

Since MRTX1133 was the first non-covalent KRAS inhibitor, we examined its activity against different RAS alleles in greater details to better understand its mechanism of action. We first tested the activity of MRTX1133 in a panel of human cancer cell lines harboring different hot-spot mutations in KRAS. Consistent with the initial reports^9,10^, two KRAS G12D mutant pancreatic cancer cell lines, Panc10.05 and AsPc-1, exhibited exquisite sensitivity to MRTX1133 (**Figure S1A**). As a control, MRTX1133 at concentration up to 1 μM showed no toxicity in the BRAF V600E mutant colorectal cancer cell line HT29, which is driven by the *BRAF* oncogene and is therefore not dependent on mutant KRAS. At 10 μM of MRTX1133, toxicity was apparent in HT29 cells (**Figure S1B**). We therefore concluded that the activity of MRTX1133 in these assays is likely to be specific at 1 μM, but not at 10 μM. Cancer cells with other *KRAS* mutant alleles, including the lung cancer cell line H358 (KRAS G12C) and the colorectal cancer cell line SW620 (KRAS G12V) also showed partial sensitivity to MRTX1133 at 1μM (**Figure S1C**). The sensitivity of these cell lines to MRTX1133 as measured by cell viability was also corroborated at the molecular level using ERK kinase phosphorylation as a readout for Ras/MAPK pathway activity. In sensitive cell lines, MRTX1133 caused a corresponding decrease in the level of phospho-ERK (pERK) (**Figure S1D**). These results suggest that the D12 mutant selectivity handle may not be essential for MRTX1133 binding to KRAS, and this compound could have inhibitory activities, albeit less potent, towards other KRAS mutants. Analogous mutation at the G12D residue also occurs in NRAS, particularly in the context of multiple myelomas (MM). We therefore tested a MM cell line that harbors a NRAS G12D mutation, INA6, for its sensitivity towards MRTX1133. Unexpectedly, INA6 cells were completely insensitive to MRTX1133 despite being highly sensitive to the MEK kinase inhibitor trametinib (**Figure S1E, S1F**). This result indicates that MRTX1133 may not inhibit NRAS G12D with similar potency as it does for KRAS G12D.

Human cancer cell lines suffer two shortcomings when used to evaluate the selectivity of a KRAS inhibitor. First, they are not isogenic, and their sensitivity to KRAS inhibition could be influenced by other co-existing mutations in the cell. Second, in addition to the mutant *KRAS* allele, these cells often retain a wild type (WT) *KRAS* allele, and they also express WT HRAS and NRAS proteins that are two other paralogs in the RAS GTPase family. Thus, a cell line’s sensitivity to MRTX1133 might result from the aggregate sensitivity of all forms of RAS protein that are expressed in the cell. To overcome these limitations, we examined the activity of MRTX1133 in a panel of isogenic “RASless” mouse embryonic fibroblasts (MEFs) that were reconstituted to express a single RAS allele. RASless MEFs were previously generated by genetically deleting all three mouse *Kras, Hras* and *Nras* genes, resulting in growth arrest in these cells^11^. The NCI RAS Initiative used lentiviral transduction to stably re-express a single human RAS cDNA in these RASless MEFs, and this was sufficient to rescue their proliferation. In these cells, proliferation is therefore strictly dependent on the sole RAS allele that was expressed (https://www.cancer.gov/research/key-initiatives/ras/ras-central/blog/2017/rasless-mefs-drug-screens). This isogenic system therefore enables us to study a single RAS allele without interference from other RAS alleles, and they are particularly useful for evaluating the selectivity of an experimental RAS inhibitor.

We first tested the sensitivity of RASless MEFs expressing different human KRAS mutants to MRTX1133. Similar to the KRAS G12D mutant cancer cell lines, RASless MEFs expressing KRAS G12D were highly sensitive to MRTX1133 with an IC_50_ of 0.07 μM (**Figure 1A**). As a control, we used RASless MEFs with a mutant BRAF V600E to further evaluate the off-target toxicity of MRTX1133 in cell-based assays. Since these cells do not express any RAS protein and their proliferation was driven by the BRAF oncoprotein, any toxicity from MRTX1133 would be strictly off-target. Consistent with the BRAF mutant cancer cell line, MRTX1133 had no impact on cell viability in the BRAF V600E RASless MEFs at concentration up to 2 μM, whereas non-specific toxicity was evident at 10 μM (**Figure 1A**). Thus, all sub-sequent studies were done at drug concentrations below 2 μM. In addition to KRAS G12D, we observed that RASless MEFs expressing KRAS G12C and KRAS G12V mutants exhibited partial sensitivity at 2 μM, whereas those expressing the KRAS G12R mutant were resistant (**Figure 1B**).

**Figure 1.**
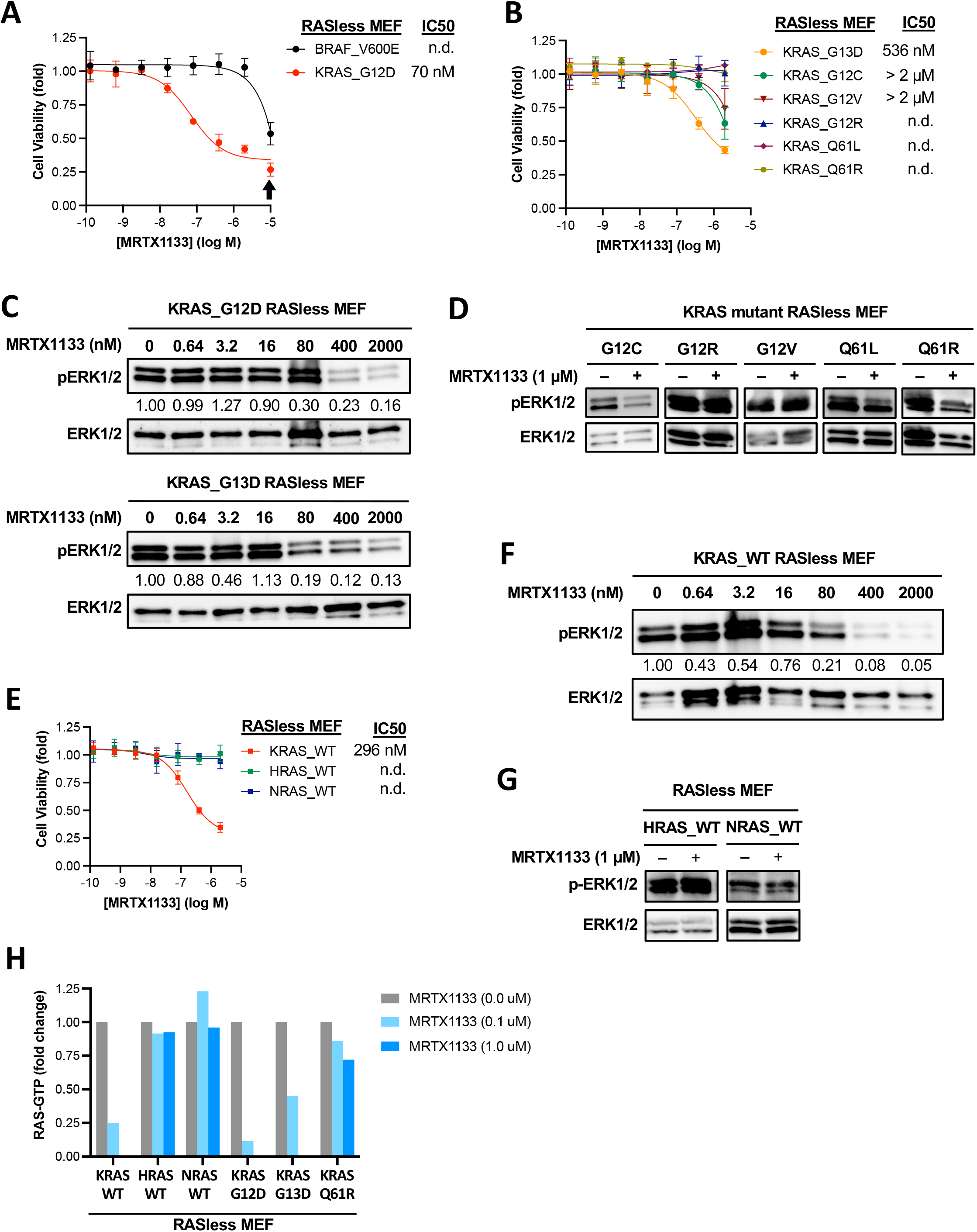
The selectivity of MRTX1133 in RASless MEFs expressing different RAS alleles. **A**. Isogenic RASless MEFs expressing human KRAS G12D or BRAF V600E were treated with MRTX1133 for 3 days. Cell viability was determined using the CellTiter-Glo assay. IC_50_ values was estimated from curve fitting from three independent repeats (error bars represent S.D.; n.d., not determined). Arrow indicates concentration of MRTX1133 (10 μM) that causes non-specific toxicity. **B**. Cell viability dose-response curves of isogenic RASless MEFs expressing different KRAS hotspot mutants. Experiments were performed analogously to those in panel **A**. **C**. RASless MEFs expressing human KRAS G12D (top panels) or KRAS G13D (bottom panels) were treated with different concentrations of MRTX1133 for one day. Cell lysates were harvested and immunoblotted for phosphorylated ERK (pERK) and total ERK. Changes in the level of pERK relative to untreated sample was quantified and shown under the pERK blots. **D**. Rasless MEFs expressing human KRAS G12C, G12R, G12V, Q61L, or Q61R mutants were treated with 1 μM MRTX1133 (+) or DMSO (-) for one day. Cell lysates were harvested and immunoblotted for pERK and total ERK. **E**. Cell viability dose-response curves of isogenic RASless MEFs expressing human WT KRAS, HRAS or NRAS protein. Experiments were performed analogously to those in panel **A**. **F**. RASless MEFs expressing human WT KRAS were treated with different concentrations of MRTX1133 for one day and the level of pERK was analyzed by immunoblotting. **G**. RASless MEFs expressing human WT HRAS or NRAS protein were treated with or without 1 μM MRTX1133 for one day and the level of pERK was analyzed by immunoblotting. **H**. Rasless MEFs expressing different wild-type or mutant RAS alleles were treated with MRTX1133 for one day. Cell lysates were harvested and the level of GTP-bound RAS protein was measured by G-LISA assay. Background-corrected signals were normalized against non-treated controls within each cell line. Data represent average of two independent repeats (the RAS-GTP level in KRAS G12D, KRAS G13D and KRAS WT cells treated with 1 μM MRTX1133 was below detection sensitivity).

Interestingly, RASless MEFs expressing the KRAS G13D mutant also showed sensitivity to MRTX1133 whereas cells expressing the KRAS Q61L and Q61R mutant were insensitive (**Figure 1B**). Concordant with cell viability data, MRTX1133 decreased pERK levels in RASless MEFs expressing G12D, G12C, G12V and G13D KRAS mutants, whereas in RASless MEFs expressing G12R, Q61L and Q61R KRAS mutants MRTX1133 had minimal impact on pERK levels (**Figure 1C, 1D**).

Next, we tested the sensitivity RASless MEFs expressing WT KRAS, HRAS or NRAS proteins. Unexpectedly, RASless MEFs expressing WT KRAS were also sensitive to MRTX1133, albeit the IC_50_ was approximately 4-fold higher than that of KRAS G12D (**Figure 1E**). This suggests that, although the D12 residue on mutant KRAS is a critical selectivity handle for MRTX1133, the scaffold of MRTX1133 has sufficient affinity for the KRAS switch II pocket such that it can still bind to KRAS when the interaction between the D12 residue on KRAS and the bicyclic piperazine ring in MRTX1122 is absent. In contrast to WT KRAS, RASless MEFs expressing WT HRAS or WT NRAS were completely insensitive to MRTX1133 (**Figure 1E**). Consistent with the cell viability result, MRTX1133 caused a dose-dependent decrease in Perk level in RASless MEFs expressing WT KRAS, whereas in RASless MEFs expressing WT HRAS or NRAS, pERK level was unchanged by MRTX1133 (**Figure 1F, 1G**).

To directly evaluate the impact of MRTX1133 on RAS protein activity, we used a G-LISA assay to measure the level of GTP-bound RAS in RASless MEFs expressing different RAS alleles following MRTX1133 treatment. Consistent with the cell viability and pERK data, we found that MRTX1133 was most effective at reducing the level of GTP-bound KRAS G12D and KRAS WT protein in the cell. It also reduced the level of GTP-bound KRAS G13D protein to a less extend. In contrast, MRTX1133 had little impact on the level of GTP-bound KRAS Q61R, or that of WT HRAS and WT NRAS protein (**Figure 1H**). These results in isogenic RASless MEFs further support the notion that MRTX1133’s activity is not solely confined to the KRAS G12D mutant, and it can inhibit WT KRAS as well as other KRAS mutants including G13D and G12C, albeit at lower potency.

The striking difference in the selectivity of MRTX1133 for KRAS over HRAS and NRAS prompted us to further investigate the mechanism that underlies its selectivity. Prior structural studies on MRTX1133 showed that its scaffold made multiple contacts throughout the G-domain of KRAS, and it binds to the GDP-bound (PDB 7RPZ) and GppCp-bound (PDB 7T47) forms of KRAS in a similar configuration^10^. We hypothesized that the paralog selectivity of MRTX1133 towards KRAS over HRAS and NRAS could be attributed to its differential interaction with the G domain of these RAS proteins. Of the amino acid residues that directly interact with MRTX1133 in co-crystal structure with KRAS G12D, all are conserved among KRAS, HRAS and NRAS proteins except histidine 95 (H95) (**Figure S2A**). In the co-crystal structure of MRTX1133 bound to KRAS G12D, the imidazole side chain of H95 makes at least three important contributions to drug binding: it forms a direct hydrogen bond with the pyrimidine ring in MRTX1133; it forms a tripartite hydrogen bonding interaction with the phenol hydroxyl group on tyrosine 64 (Y64) and the pyrimidine ring in MRTX1133; and it forms a T-shaped interaction with the phenol ring of Y96, stabilizing the hydrogen bond between Y96 and the pyrimidine ring on MRTX1133 (**Figure S2B**). In HRAS and NRAS, the corresponding residues are glutamine (Q95) and leucine (L95), respectively (**Figure S2A**). Thus, it is plausible to postulate that the non-conserved residue H95 on KRAS provides a critical selectivity handle for MRTX1133, and substitution of H95 with Q and L, as seen in HRAS and NRAS, respectively, could substantially disrupt MRTX1133 binding.

To test this hypothesis, we generated reciprocal mutations in G12D mutant KRAS, HRAS and NRAS where H95 in KRAS was mutated to Q95 and L95 as in HRAS and NRAS, respectively, and Q95 in HRAS and L95 in NRAS was mutated to H95 as in KRAS. We stably expressed the cDNA of these double mutants in the context of the KRAS G12D RASless MEFs so we could directly compare their activity against the sensitive KRAS G12D mutant in the same cells (**Figure S2C**). An N-terminal HA-tag was used to distinguish the double mutants from the existing KRAS G12D allele. In support of our hypothesis, expression of the HA-KRAS G12D/H95Q and the HA-KRAS G12D/H95L double mutants rendered cells completely resistant to MRTX1133 (**Figure 2A**). Using RAS-GTP pulldown assay to directly measure the impact of MRTX1133 on these double mutants, we found that MRTX1133 was ineffective at reducing the level of GTP-bound form of either double mutant compared to KRAS G12D (**Figure 2B**). Consequently, expression of these double mutants also prevented the downregulation of pERK by MRTX1133 (**Figure 2C**).

**Figure 2.**
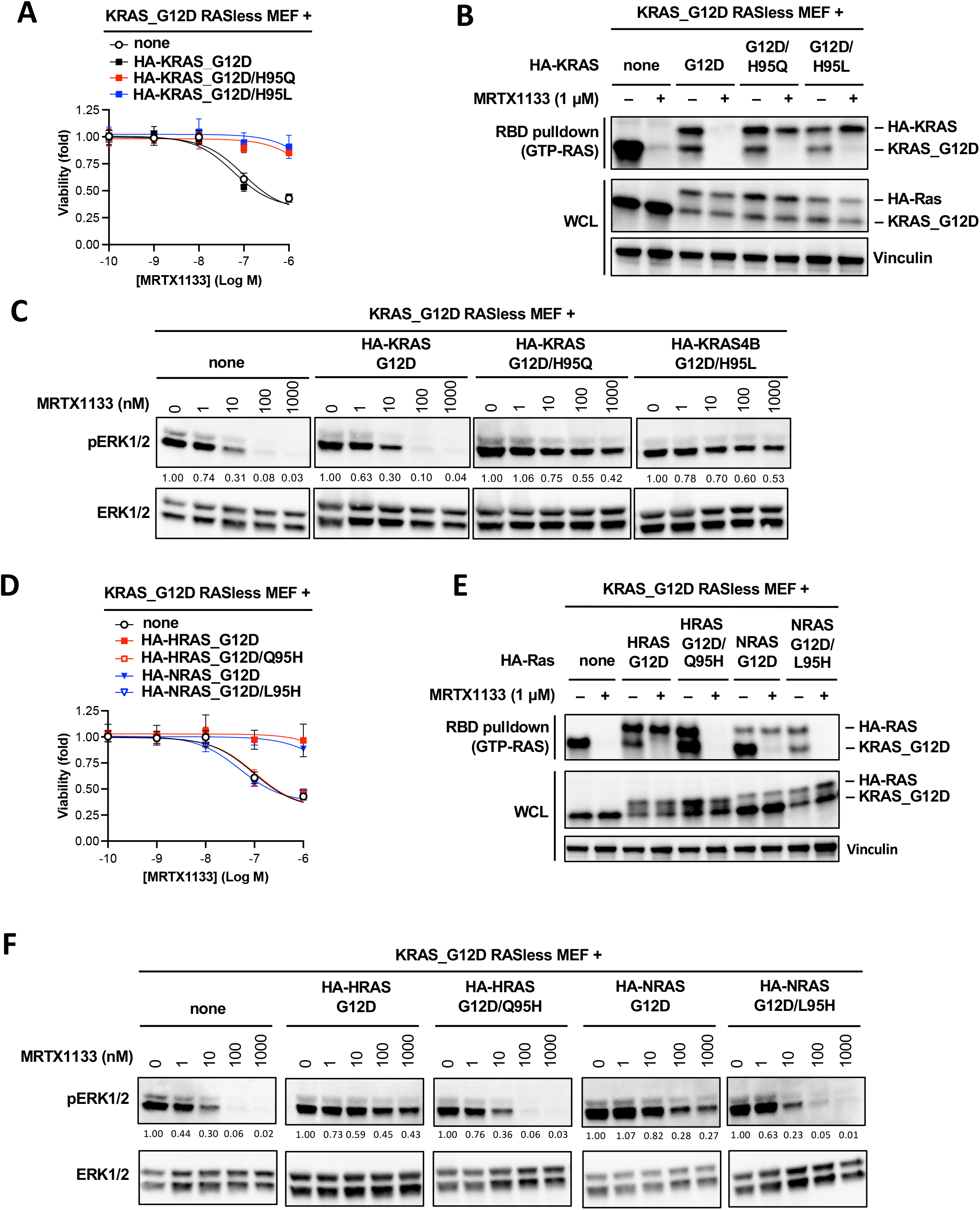
Histidine 95 on KRAS determines the paralog selectivity of MRTX1133. **A**. KRAS G12D RASless MEFs were stably transduced with cDNAs expressing HA-tagged KRAS G12D, KRAS G12D/H95Q or KRAS G12D/H95L. Resulting cells were treated with MRTX1133 for 3 days. Cell viability was determined using the CellTiter-Glo assay. Data shown as average of three independent repeats (error bars represent S.D). **B**. Cell lines used in panel **A** were treated with 1 μM MRTX1133 (+) or DMSO (-) for one day. Cell lysates were harvested and GTP-bound KRAS levels was measured using RBD pulldown assay (top panel). Same lysates were immunoblotted to determine the input levels of RAS proteins. Untagged KRAS G12D and HA-tagged KRAS were detected with a pan-Ras antibody. **C**. Cell lines used in panel **A** were treated with MRTX1133 for one day. Cell lysates were immunoblotted for pERK and total ERK. Changes in the level of pERK relative to untreated sample was quantified and shown under the pERK blot. **D-F**. The same experiments as in panels A-C were performed in KRAS G12D RASless MEFs transduced with cDNAs expressing HA-tagged HRAS G12D, HRAS G12D/Q95H, NRAS G12D, or NRAS G12D/L95H mutants.

Consistent with the notion that MRTX1133 is unable to inhibit HRAS and NRAS G12D protein, expression of HA-HRAS G12D and HA-NRAS G12D single mutant rendered cells completely resistant to MRTX1133. In contrast, cells expressing the HA-HRAS G12D/Q95H and the HA-NRAS G12D/L95H double mutants remain fully sensitive to MRTX1133 (**Figure 2D**). These sensitivity profiles were corroborated by the impact of MRTX1133 on the level of GTP-bound Ras: the level of GTP-bound HA-HRAS G12D and HA-NRAS G12D single mutants remained unchanged in response to MRTX1133, whereas the level of GTP-bound HA-HRAS G12D/Q95H and the HA-NRAS G12D/L95H double mutants was effectively reduced by MRTX1133 to a similar extent as KRAS G12D (**Figure 2E**). This differential sensitivity was also reflected by the effect of MRTX1133 on pERK level in these cells (**Figure 2F**). Together, these results indicate that the non-conserved H95 residue on KRAS presents an essential selectivity handle that drives the selectivity of MRTX1133 towards KRAS over HRAS and NRAS. The inability of MRTX1133 to inhibit the G12D mutant of HRAS and NRAS is likely due to the presence of Q95 and L95 on these paralogs, respectively, that interfere with high affinity binding by MRTX1133.

Our analysis of the prototypical KRAS G12D inhibitor MRTX1133 led to several important insights that have impact for the future development of this class of non-covalent KRAS inhibitors. First, the D12 mutant residue is critical, but not essential, for inhibitor activity. Our data showed that MRTX1133 can clearly inhibit WT KRAS, albeit at lower potency. In retrospect, this was perhaps not entirely surprising. Although the bicyclic piperazine ring on MRTX1133 was optimized for binding to the D12 residue, to gain sufficient affinity without the advantage of crosslinking, the MRTX1133 scaffold was also extensively optimized to bind to the switch II pocket on KRAS. Thus, losing the hydrogen bond interaction between the piperazine ring with the D12 residue seems to reduce, but not abolish, its binding to KRAS. This explains the partial activity of MRTX1133 towards other KRAS mutants including G12C and G13D and towards WT KRAS. The activity of MRTX1133 towards WT KRAS could also be partially attributed to its preference for binding to the GDP-bound form of KRAS^10^. As WT KRAS protein cycles much more rapidly through the GDP-bound state than mutant KRAS, it might be more amenable for MRTX1133 binding. In support of this notion, although the Q61 side chain on KRAS does not directly interact with MRTX1133, both Q61R and Q61L mutants of KRAS were resistant to MRTX1133 in our assay, likely due to the ability of these mutants to remain in the GTP bound state for an extended period of time^12^.

Second, we showed that MRTX1133 exhibits exquisite selectivity for KRAS over its two closely related paralogs HRAS and NRAS. Our reciprocal mutation experiment demonstrated that this is entirely due to the non-conserved H95 residue on KRAS which seems to play an essential role in stabilizing the binding of MRTX1133. Given its activity towards WT KRAS protein, the wide difference in sensitivity towards MRTX1133 in human cancer cell lines with and without KRAS G12D mutation^10^ is likely due, at least in part, to its ability to spare HRAS and NRAS protein. Thus, the H95 selectivity handle on KRAS could be critical for reducing the on-target toxicity of this inhibitor in normal tissues. It is likely that a pan-Ras inhibitor would be too toxic to be useful in the clinic. However, blocking mutant KRAS signaling while sparing physiological HRAS and NRAS signaling could substantially reduce the sensitivity of normal tissues that are not addictive to the *KRAS G12D* oncogene towards MRTX1133. The KRAS G12C inhibitor sotorasib has a distinct scaffold compared to MRTX849 and MRTX1133. In the co-crystal structure of KRAS G12C with sotorasib, all the amino acids that directly interact with sotorasib are conserved across the three RAS paralogs, and the non-conserved H95 does not directly interact with sotorasib^5^. Thus, it is reasonable to postulate that a KRAS G12D inhibitor based on the sotorasib scaffold might also inhibit HRAS G12D and NRAS G12D proteins and lack paralog selectivity.

Third, our study suggests a resistance mechanism to MRTX1133 that will likely be encountered in clinical studies. MRTX1133 was evolved from, and shares structural similarity with, the KRAS G12C inhibitor adagrasib. In patients treated with adagrasib, one form of acquired resistance mechanism is secondary mutation in residues H95 and Y96 on KRAS that disrupts adagrasib binding^13,14^. Adagrasib and MRTX1133 have related heterocyclic core scaffolds. In the co-crystal structure of adagrasib covalently bound to KRAS G12C, H95 and Y96 form hydrogen bonds with the ligand pyrimidine ring^4^ (**Figure S2D**). Our data showed that H95 is critical for the non-covalent binding of MRTX1133 to KRAS. We therefore anticipate that secondary mutation at H95 will drive clinical resistance to MRTX1133. Since Y96 forms a hydrogen bond with the pyrimidine ring in MRTX1133, we reasoned that Y96 mutation could also disrupt MRTX1133 binding. Indeed, in KRAS G12D RASless MEFs, the additional expression of a HA-KRAS G12D/Y96D double mutant render cells fully resistant to MRTX1133 (**Figure S3A**), and the GTP-bound level of this double mutant remained unchanged in the presence of the drug (**Figure S3B**). Consequently, MRTX1133 was unable to effectively reduce pERK level in cells expressing the HA-KRAS G12D/Y96D double mutant (**Figure S3C**). Furthermore, in the KRAS G12D mutant human pancreatic cancer cell line Panc10.05, which is highly sensitive to MRTX1133 (**Figure S1A**), expression of the KRAS G12D/H95Q, KRAS G12D/H95L and KRAS G12D/Y96D double mutants all rendered the cells fully resistant to MRTX1133 (**Figure S3D, S3E**). Thus, analogous to adagrasib, secondary mutation at the H95 and Y96 residues on KRAS will likely be a significant mechanism of acquired resistance to MRTX1133 and its derivatives in clinical studies.

Lastly, our study suggests that “pan-KRAS” inhibitors with broader spectrum of activities could be developed by exploiting the KRAS H95 selectivity handle. Our characterization of the selectivity profile of MRTX1133 indicates that optimizing binding for a mutant residue, while highly desirable, may not be necessary for the development of a high-affinity inhibitor of KRAS. It might be possible to develop a chemically related scaffold that preserves binding to H95 but does not extend interaction towards the G12 or G13 position on KRAS. Given that the three RAS paralogs tend to be co-expressed in many adult tissues, a pan-KRAS inhibitor that blocks both mutant and WT KRAS while sparing HRAS and NRAS could be highly toxic in KRAS addicted cancer cells yet well tolerated in normal tissues^15^. Conversely, altering the scaffold such that it specifically interacts with Q95 or L95 but not with H95 could aid the development of HRAS and NRAS paralog-selective inhibitors, respectively, to target HRAS and NRAS mutant tumors.

## Materials and Methods

### Cell culture and reagents

Human cancer cell lines were grown in RPMI-1640 medium (Lonza #12-702F) supplemented with 10% fetal bovine serum (Thermo Fisher Scientific #10438026) and 100 units/mL of penicillin and 100 ug/mL of streptomycin (Thermo Fisher #15140). RaslessMEF cells expressing different RAS alleles were generated by and obtained from the National Cancer Institute (NCI) RAS Initiative (https://www.cancer.gov/research/key-initiatives/ras/ras-central/blog/2017/rasless-mefs-drug-screens), and were grown in DMEM (Lonza #12-604F) supplement with 10% fetal bovine serum (Thermo Fisher Scientific #10438026) and 100 units/mL of penicillin and 100 ug/mL of streptomycin (Thermo Fisher #15140). Puromycin (Thermo Fisher Scientific #A1113803) was added in the culture media to a final concentration of 2.5 μg/ml to maintain the stable expression of wild-type HRAS in RaslessMEF cells. Blasticidin (Thermo Fisher Scientific #A1113903) was added in the culture media to a final concentration of 4 μg/ml to maintain the stable expression of all the other wild-type or mutant RAS. All the cells were cultured at 37 °C in a humidified 5% CO2 incubator. MRTX1133 (ChemGood #C-1420) and Trametinib (Selleckchem #S2673) were dissolved in DMSO at the stock concentration of 10 mM.

### Cell viability assay

Cell viability assay was performed using CellTiter-Glo One reagent (Promega #G8462) following the manufacturer’s instructions. Briefly, human cancer cells (2000 cells per well) or Rasless MEF cells (3000 cells per well) were plated and cultured in black-walled clear bottom 96-well tissue culture-treated plates (Corning #3603) for 24 hours. Next day, equal volume of media containing 2X MRTX1133 or trametinib was added into wells such that the final concentrations of the drug reached the desired concentration in culture. Cells were treated for 3 days, and CellTiter-Glo assay was performed at the end point. Luminescence signal was detected on a TECAN Infinite M200 plate reader (Tecan Trading AG, Switzerland), and cell viability was calculated by normalizing the luminescence signal to non-treated conditions. Dose-response curve was fitted and plotted using GraphPad Prism 9.0 (GraphPad Software, LLC).

### Immunoblotting and antibodies

To collect whole cell extracts for immunoblotting, cells were kept on ice and washed with ice-cold DPBS (Corning #21-031-CV) twice then lysed in RIPA lysis buffer (50 mM Tris-HCl pH7.4, 150 mM NaCl, 1 mM EDTA, 1% NP-40, 0.25% sodium deoxycholate, 2 mM Na_3_VO_4_, 20 mM sodium pyrophosphate, 20 mM NaF, 20 mM β-glycerophosphate) containing additional protease inhibitors (Sigma-Aldrich #11836170001) and phosphatase inhibitors (Sigma-Aldrich # 4906837001). Total protein concentration was determined by BCA assay (Thermo Fisher Scientific #23225). Cell lysate was mixed with Laemmli Sample Buffer (Bio-Rad #1610747) and denatured at 95°C for 10 minutes. Samples were separated by SDS-PAGE in an 8-16% gradient Tris/Glycine gel (Bio-Rad #5671105) then transferred onto a nitrocellulose membrane by using the Trans-Blot Turbo RTA transfer kit (Bio-Rad #1704271). To detect protein abundance, membranes were first blocked in 5% blocking milk (Bio-Rad #1706404) in TBS (Quality Biological #351-086-131) containing 0.1% Tween-20 (Sigma #P1379) for 30 minutes, then hybridized with primary antibodies (Supplemental Table 1) following manufacturers’ instructions. After primary antibody hybridization, membranes were washed five minutes in TBS containing 0.1% Tween-20 for three times then hybridized with horseradish peroxidase (HRP)-conjugated secondary antibodies (Supplementary Table 1) in TBS containing 0.1% Tween-20 for one hour. After secondary antibody hybridization, membranes were washed 10 minutes in TBS containing 0.1% Tween-20 for three times. HRP signal was detected by using the Immobilon Forte Western HRP Substrate solution (Sigma #WBLUF0100) and acquired by using a ChemiDoc Touch Imaging System (Bio-Rad Laboratories, Inc). Chemiluminescence signal was quantified using the Image Lab Software (Bio-Rad Laboratories, Inc).

### Construction of mutant RAS expression vectors

Plasmids harboring WT and G12D mutant KRAS, HRAS, and NRAS cDNA were obtained from the NCI RAS Initiative. RAS double mutants with point-mutations in the 95^th^ and 96^th^ amino acid residues were generated by site-directed mutagenesis (Agilent Technologies #200521) using the primers listed in Supplementary Table 2 per manufacturer’s recommended PCR conditions. The coding region of the RAS cDNA and point mutations were verified by Sanger Sequencing. To construct N-terminal HA-tagged RAS-expressing plasmids, the coding region of RAS cDNA was first PCR-amplified from the plasmids above by using CloneAmp HiFi PCR Premix (Takara #639298) and the primers listed in Supplementary Table 3 using manufacturer’s recommended PCR conditions. The PCR products were purified by gel purification (Qiagen #28706) then digested with the restriction enzymes NotI (New England Biolabs #R3189) and BamHI (New England Biolabs #R3136). Digested PCR fragments were gel-purified (Qiagen #28104) and ligated into the pLVX-Puro vector (Takara #632164) between the PspOMI and BamHI restriction sites by using T4 DNA ligase (New England Biolabs #M0202). Ligated plasmids were transformed into NEB stable cells (New England Biolabs #C3040) following manufacturer’s instructions, and single colonies were isolated and grown for plasmid preparation (Qiagen #28115). The entire coding region of the HA-RAS cDNA were verified by Sanger Sequencing.

### Lentivirus production and generation of HA-RAS-expressing stable cells

To produce lentivirus, 293FT cells were co-transfected with 1 μg of the pLVX-Puro vectors carrying the cDNAs expressing HA-tagged RAS variants, 0.5 μg of pMD2.G (Addgene #12259), 0.75 μg of psPAX2 (Addgene #12260) in 6-well plates by using TransIT-293 transfection reagent (Mirus #MIR2705) following manufacturer’s instructions. Sixty hours after transfection, virus-containing media was collected, aliquots, and frozen at −80°C. To generate cell lines stably expressing HA-RAS variants, RaslessMEF cells and Panc10.05 cells were plated and grown overnight. Culture medium was replaced with lentivirus-containing medium at the ratio of 100 μL virus-containing medium per 25,000 cells seeded in the presence of 6 μg/mL polybrene (Thermo Fisher Scientific #TR1003G), followed by centrifugation at 2,000 rpm in room temperature for 30 minutes. Cells were cultured and incubated with the lentiviral media at 37°C in a humidified 5% CO2 incubator for 24 hours, and then cultured in normal culture medium for another 24 hours. Cells stably expressing the transduced genes were selected with 5 μg/ml puromycin for 3 days and subsequently maintained in medium containing 2.5 μg/ml puromycin. HA-RAS expression was verified by immunoblotting.

### RAS-GTP assays

To determine the inhibitory effect of MRTX1133 on RAS GTP loading in RaslessMEF cells expressing a single allele of RAS variant, the colorimetric-based RAS G-LISA activation assay (Cytoskeleton #BK131) was used. Briefly, cells were treated with MRTX1133 or DMSO control for 3 days, and cell lysates were collected on ice, aliquoted, and frozen in liquid nitrogen. Total protein concentration in the lysates were determined by using the Pierce BCA Protein Assay Kit (Thermo Fisher Scientific #23225). Briefly, 25 μg of totally lysates were subjected to G-LISA assay in duplicated wells following the manufacturer’s instructions, and another 15 ug of total lysates were subjected to immunoblotting to determine RAS expression level as the input. Colorimetric signal in each well was quantified using a TECAN plate reader, and a background signal from wells containing no lysate was subtracted to yield a specific signal. If the signal of a sample well was below that of the back background signal, the specific signal of that well was treated as being zero (i.e. below detection limit). Relative RAS-GTP level was calculated by normalizing against DMSO-treated wells and adjusted by the input level from the corresponding immunoblot.

To determine the inhibitory effect of MRTX1133 on HA-RAS with double mutations, the RAS-GTP pull-down assay kit (Cytoskeleton #BK008) was used. Briefly, cells stably expressing HA-RAS mutant was treated with MRTX1133 or DMSO control for 24 hours. Cell lysate was collected on ice, aliquoted, and frozen in liquid nitrogen. Total protein concentration in the lysate was determined with BCA assay, and 300 μg of totally lysate was subjected to RBD RAS-GTP pull-down assay following the manufacturer’s instruction. Ras protein level in the pull down was analyzed by by immunoblotting using RAS antibody (Sigma-Aldrich #OP40). Total was lysate was also subjected to immunoblotting to determine RAS expression level as the input.

## Acknowledgements

We would like to thank the National Cancer Institute RAS Initiative for generously providing the panel of Rasless MEF cell lines and human RAS cDNA plasmids for cloning, the CCR Genomics Core facility for sequencing support, and Dr. John S Schneekloth at the NCI for helpful discussion.

## Funding

This research was supported by the Intramural Research Program of the National Cancer Institute, NIH.

## Author contributions

All the authors designed the experiments and approaches. M.A.K, J.J.W.H, Y-H. L, and C-S. L. performed the experiments, analyzed the results, and generated the figures. C-S. L and J. L. wrote and prepared the manuscript.

## Supplementary information

**Supplementary Table 1.**
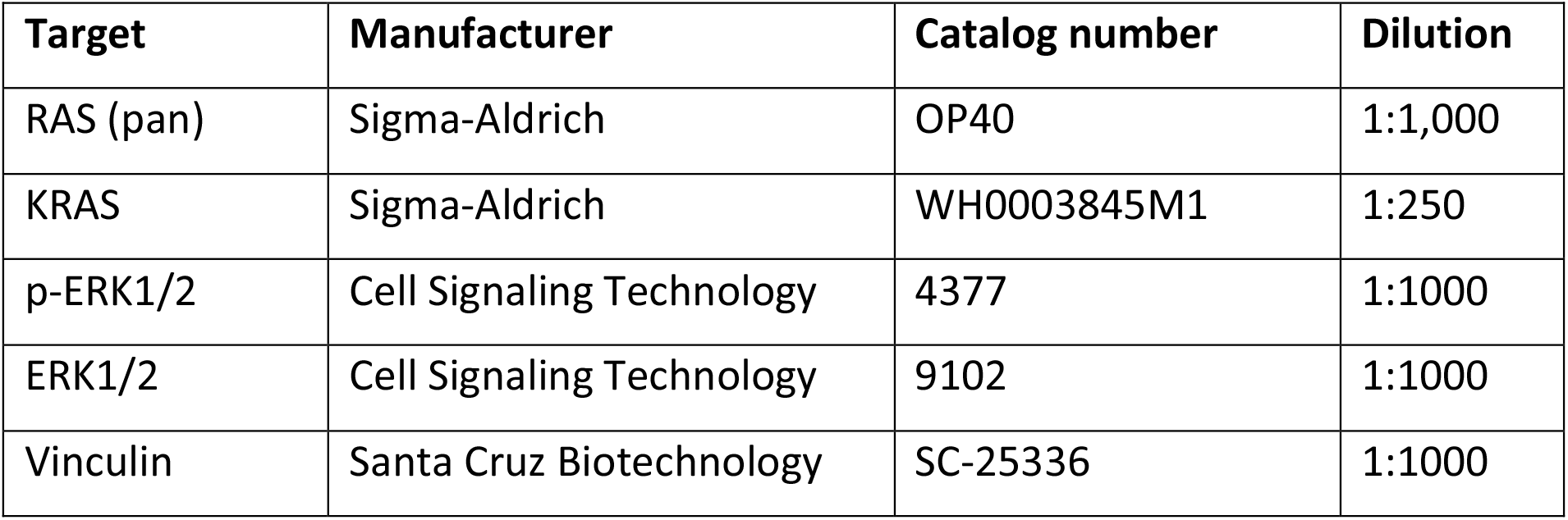
Antibodies for immunoblotting.

| Target | Manufacturer | Catalog number | Dilution |
| --- | --- | --- | --- |
| RAS (pan) | Sigma-Aldrich | OP40 | 1:1,000 |
| KRAS | Sigma-Aldrich | WH0003845M1 | 1:250 |
| p-ERK1/2 | Cell Signaling Technology | 4377 | 1:1000 |
| ERK1/2 | Cell Signaling Technology | 9102 | 1:1000 |
| Vinculin | Santa Cruz Biotechnology | SC-25336 | 1:1000 |

**Supplementary Table 2.**
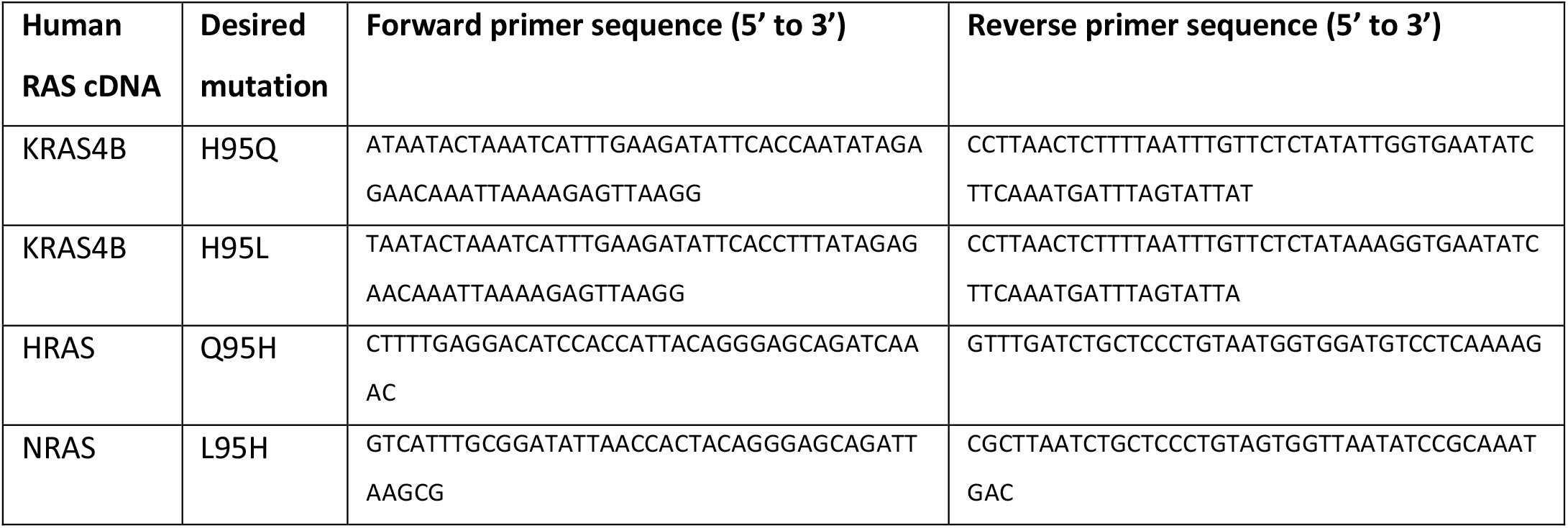
Site-directed mutagenesis primers.

**Supplementary Table 3.**
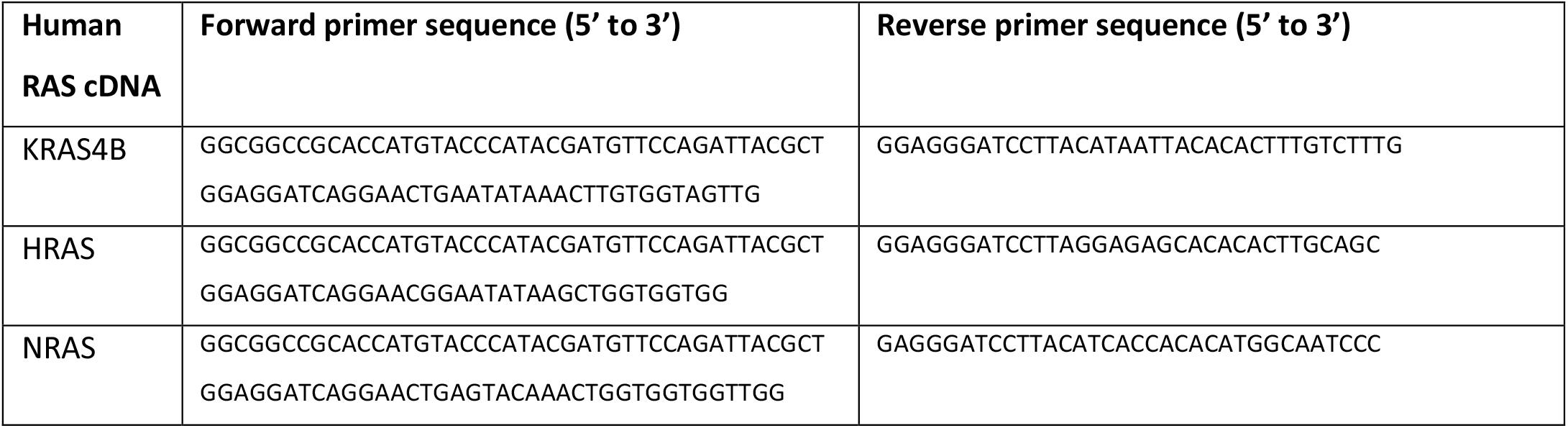
Primers for PCR-cloning HA-RAS into pLVX-Puro vector.

## Supplementary Figure Legends

**Supplementary Figure S1.**
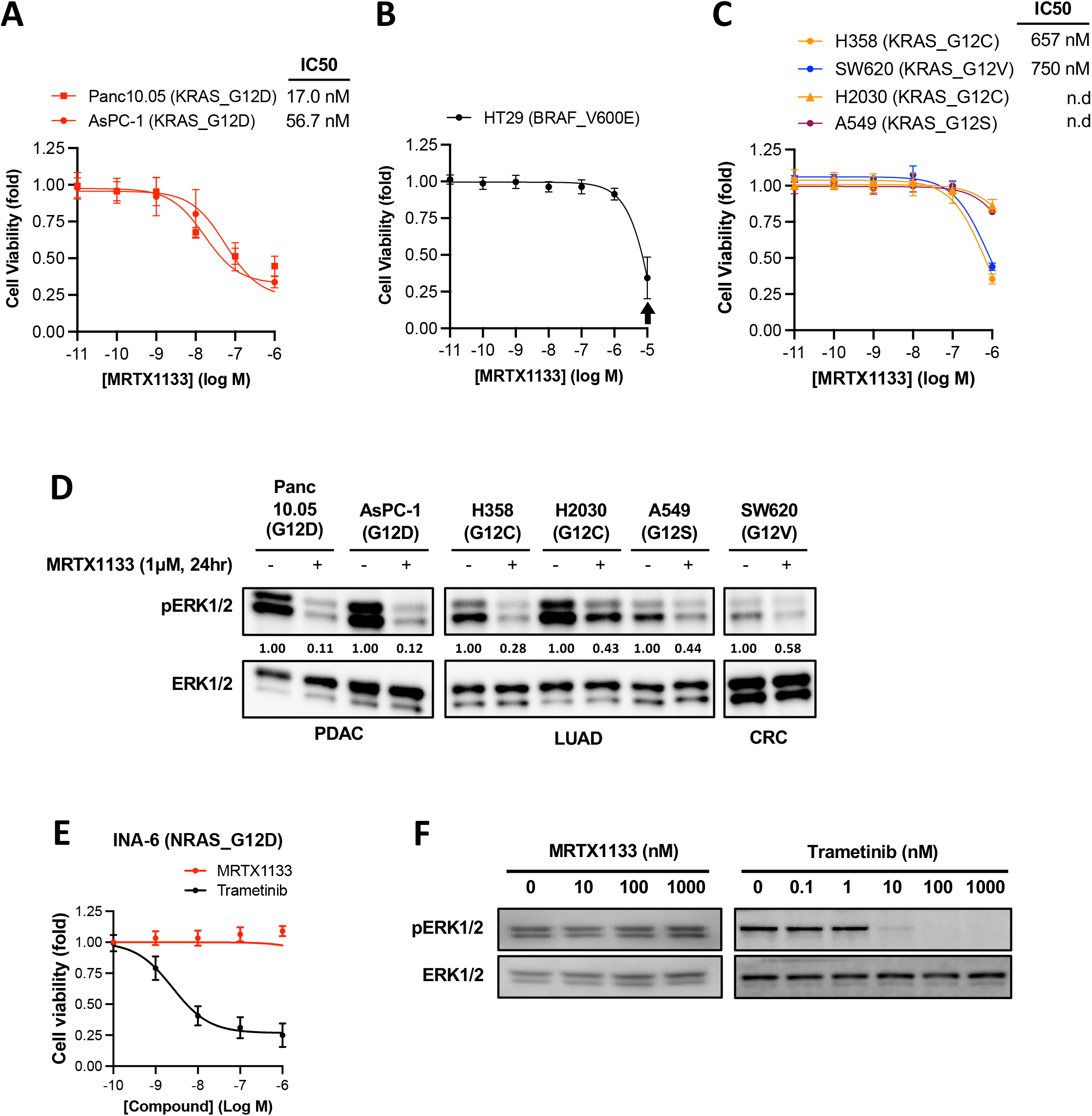
Selectivity of MRTX1133 in RAS mutant human cancer cell lines. **A-C**. Human cancer cell lines harboring different KRAS or BRAF mutations were treated with MRTX1133 for 3 days. Cell viability was determined using the CellTiter-Glo assay. IC50 values was estimated from curve fitting using three independent repeats (error bars represent S.D.; n.d., not determined). Arrow indicates concentration of MRTX1133 (10 μM) that causes non-specific toxicity. **D**. Human cancer cell lines harboring different KRAS or BRAF mutations were treated with 1 μM MRTX1133 (+) or DMSO (-) for one day. Cell lysates were harvested and immunoblotted for pERK and total ERK. Changes in the level of pERK relative to untreated sample was quantified and shown under the pERK blots. **E**. INA6 cells were treated with MRTX1133 or Trametinib for 3 days. Cell viability assay was performed analogously to that in panel **A**. **F**. INA6 cells were treated with different concentrations of MRTX1133 or trametinib for one day. Cell lysates were harvested and immunoblotted for pERK and total ERK.

**Supplementary Figure S2.**
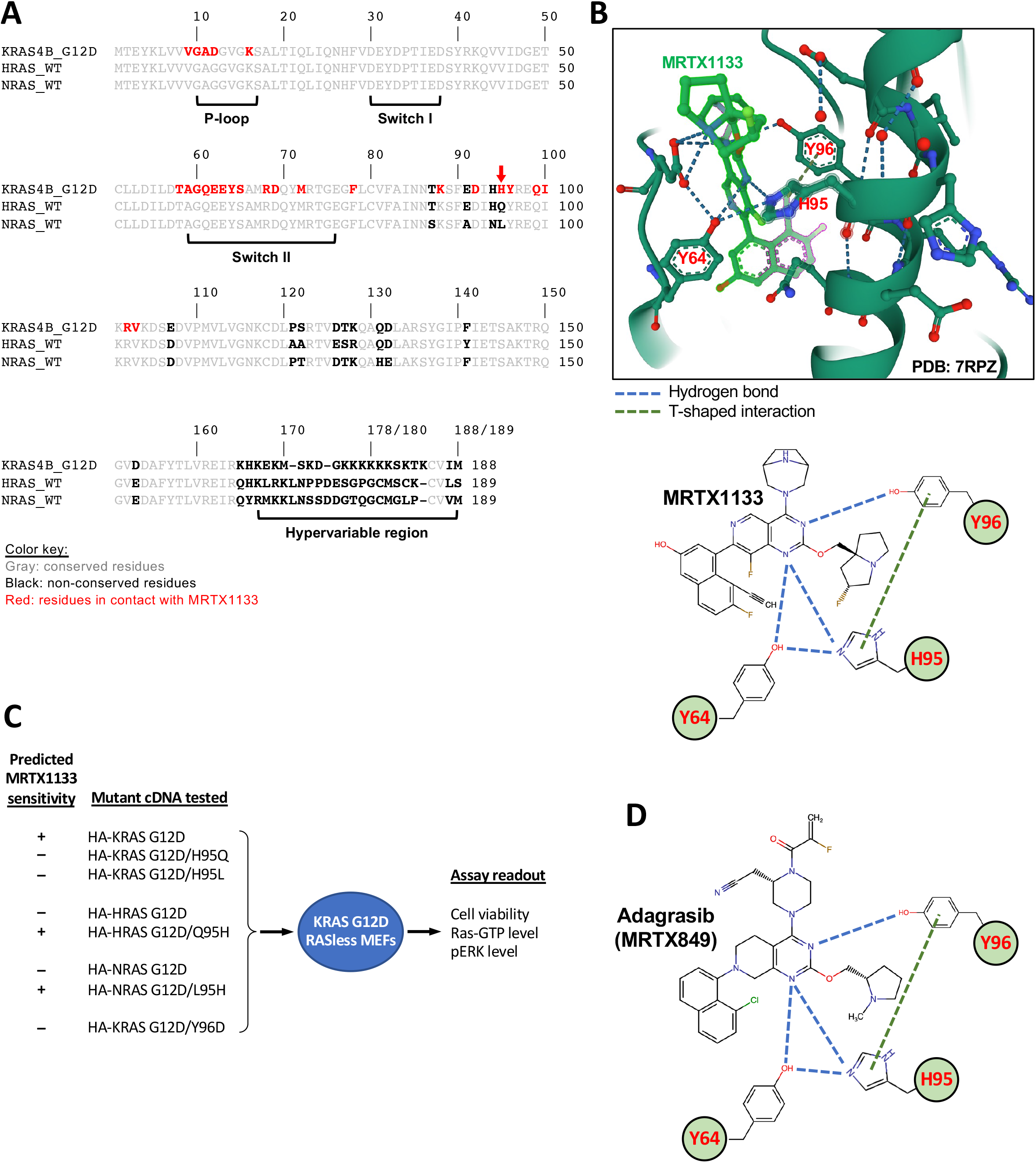
The non-conserved histidine 95 on KRAS makes critical contribution to MRTX1133 ligand binding. **A**. Amino acid sequence alignment of human KRAS4B G12D, WT HRAS, and WT NRAS. The P-loop, Switch I and Switch II domains, and the hypervariable region are labeled. Amino acid residues of KRAS G12D that interact with MRTX1133 are colored in red. Residues that are **A. A**. conserved among the three RAS sequences are colored in gray, while the residues that are not conserved are bold and colored in black. Among all the MRTX1133-interacting residues on KRAS G12D, only H95 (red arrow) is not conserved in HRAS and NRAS. **B**. Interactions of MRTX1133 with H95 on KRAS G12D. Top panel, structure of the MRTX1133 binding pocket in KRAS G12D in the GDP-bound form illustrating the role H95 plays in stabilizing ligand binding (PDB 7RPZ). Bottom panel, a schematic view of the interactions between MRTX1133 and Y64, H95, and Y96 residues of KRAS. **C**. Experimental design for testing the hypothesis that H95 on KRAS provides a critical selectivity handle for MRTX1133. Different HA-tagged double RAS mutants were transduced into KRAS G12D RASless MEFs to test how they alter the cells sensitivity to MRTX1133. **D**. A schematic view of the interactions between adagrasib and Y64, H95, and Y96 residues of KRAS (based on PDB 6UT0).

**Supplementary Figure S3.**
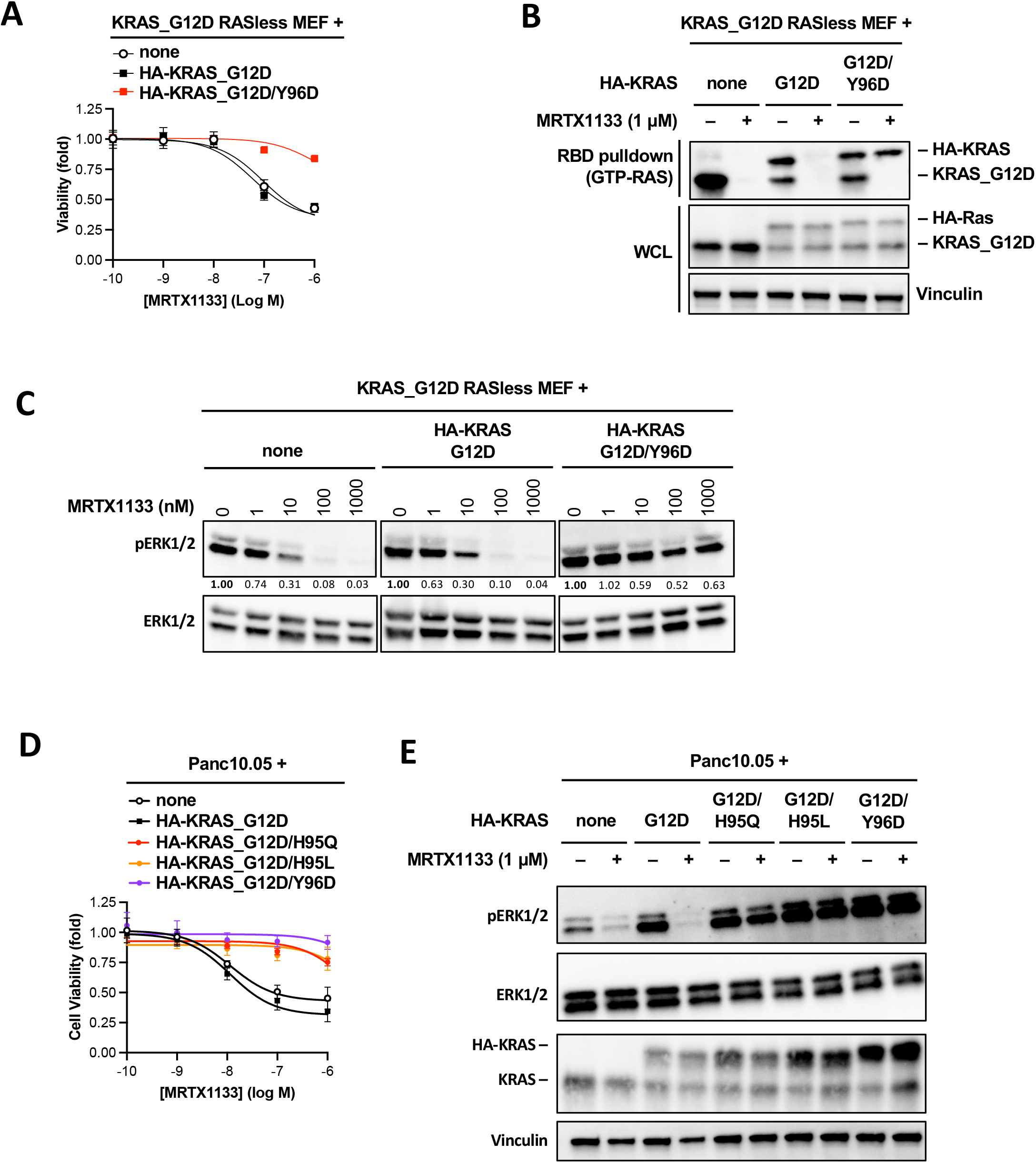
Mutation in H95 and Y96 on KRAS as a potential mechanism for acquired resistance to MRTX1133. **A**. KRAS G12D Rasless MEFs were stably transduced with HA-tagged KRAS G12D or KRAS G12D/Y96D. Cells were treated with MRTX1133 for 3 days and cell viability was determined using CellTiter-Glo assay. Dose-response curves were fitted from three independent repeats (error bars represent S.D.). **B**. Cells used in panel **A** were treated with 1 μM MRTX1133 (+) or DMSO (-) for one day. Whole cell lysates were harvested and subjected to RAS-GTP pulldown assay (top panel) to determine GTP-bound KRAS levels. Lysates were also subjected to immunoblotting to determine the input levels. **C**. Cells used in panel **A** were treated with different concentrations of MRTX1133 for one day. Cell lysates were harvested and immunoblotted for pERK and total ERK. Changes in the level of pERK relative to untreated sample was quantified and shown under the pERK blots. **D**. Human pancreatic cancer cell line Panc10.05 were stably transduced with HA-tagged KRAS mutants as indicated. Cells were treated with MRTX1133 for 3 days and cell viability was determined using CellTiter-Glo assay. Dose-response curves were fitted from three independent repeats (error bars represent S.D.). **E**, Cells used in panel **D** were treated with 1 μM MRTX1133 (+) or DMSO (-) for one day. Cell lysates were harvested and immunoblotted for pERK and total ERK.

## Notes

### Competing Interest Statement

The authors have declared no competing interest.

